# Neuronal GPCR NMUR-1 regulates energy homeostasis in response to pathogen infection

**DOI:** 10.1101/2024.07.09.602733

**Authors:** Phillip Wibisono, Yiyong Liu, Kenneth P Roberts, Dodge Baluya, Jingru Sun

## Abstract

A key question in current immunology is how the innate immune system generates high levels of specificity. Our previous study in *Caenorhabditis elegans* revealed that NMUR-1, a neuronal G protein-coupled receptor homologous to mammalian receptors for the neuropeptide neuromedin U (NMU), regulates distinct innate immune responses to different bacterial pathogens. Here, by using quantitative proteomics and functional assays, we discovered that NMUR-1 regulates F_1_F_O_ ATP synthase and ATP production in response to pathogen infection, and that such regulation contributes to NMUR-1-mediated specificity of innate immunity. We further demonstrated that ATP biosynthesis and its contribution to defense is neurally controlled by the NMUR-1 ligand CAPA-1 and its expressing neurons ASG. These findings indicate that NMUR-1 neural signaling regulates the specificity of innate immunity by controlling energy homeostasis as part of defense against pathogens. Our study provides mechanistic insights into the emerging roles of NMU signaling in immunity across animal phyla.

## INTRODUCTION

The soil dwelling nematode *Caenorhabditis elegans* is a powerful model for studying host-pathogen interactions on the whole animal scale. This small invertebrate measuring slightly over 1mm in length is susceptible to infection from multiple bacterial and viral pathogens, including some known human pathogens (1-3). Unlike vertebrates and some invertebrates, *C. elegans* does not have an adaptive immune system or professional immune cells; instead, the nematode relies on a localized innate immune response and behavior responses to defend itself (4,5). Upon infection, *C. elegans* mounts an immune defense by triggering evolutionarily conserved signaling cascades such as the mitogen-activated protein kinase (MAPK) pathway, the DAF-2/insulin-like pathway, and/or the transforming growth factor β homolog (TGF-β) pathway (6-8). Activation of these pathways promotes the expression of a variety of defense genes, allowing *C. elegans* to fight pathogen infection (4,9,10).

These defense responses must be tightly regulated as a lack of control can be harmful or even detrimental to the host. For example, animals lacking a functional NPR-1, a G protein-coupled receptor (GPCR) expressed in neurons, have decreased survival against pathogens such as *Pseudomonas aeruginosa* due to dysregulated gene expression in the p38/PMK-1 MAPK pathway. However, loss-of-function mutations in other neuronal GPCRs can be beneficial during pathogen infection, as in the case of a mutation in OCTR-1. OCTR-1 null animals have an increase in survival against *P. aeruginosa* because of an increase in stress response proteins in the unfold protein response (UPR) pathway (11-14). We recently discovered that a neuronal GPCR, neuromedin receptor 1 (NMUR-1), regulates distinct immune responses to different pathogens by controlling the expression of various defense genes at the transcriptional level (15). When challenged with Gram-negative pathogen *Salmonella enterica, nmur-1(ok1387)* knockout mutants display an enhanced survival phenotype due to upregulation of UPR related genes. However, the *nmur-1(ok1387)* mutant animals display a reduced survival phenotype when exposed to Gram-positive pathogen *Enterococcus faecalis* because of a lack of expression of C-type lectin genes. This previous research focused on changes at the transcriptome level and how NMUR-1 affects gene expression. In the current study, we investigate the role of NMUR-1 by examining how NMUR-1 regulates the host proteome during distinct pathogen challenges. We examined the proteomes of both wild-type (WT) and *nmur-1(ok1387)* mutant animals during *S. enterica* and *E. faecalis* infections and discovered that the mutant animals have a significant decrease in the abundance of proteins involved in transmembrane transport. More specifically, the downregulated proteins are involved in the movement of ions across cell membranes, which include subunits of the F_1_F_O_ ATP synthase complex, such as ATP5O, ATP5PD, and ATP5PF. F_1_F_O_ ATP synthase is a large protein complex necessary to produce ATP. The flow of a non-equilibrium proton gradient generated by the electron transport chain through the mitochondrial membrane drives the rotation of the F_1_F_O_ molecular motors to generate ATP (16,17). We found that reducing ATP production by inhibiting the F_1_F_O_ ATP synthase chemically or genetically caused WT N2 animals to mimic the survival phenotypes of *nmur-1(ok1387)* animals during either *S. enterica* or *E. faecalis* infection. Consistent with our proteomics data, functional loss of NMUR-1 reduces ATP concentrations during pathogen infection. These results indicate that NMUR-1 regulates energy homeostasis by controlling ATP generation, and that downregulation of ATP production is partially responsible for the distinct survival phenotypes of *nmur-1(ok1387)* mutant animals exposed to *S. enterica* and *E. faecalis*. Moreover, we show that both the NMUR-1 endogenous ligand CAPA-1 and its expressing neurons ASG regulate ATP biosynthesis during *E. faecalis* infection and are required for *C. elegans* survival against this pathogen, demonstrating that ATP biosynthesis and its contribution to defense is neurally controlled by the CAPA-1/NMUR-1/ASG signaling. Overall, our study revealed NMUR-1-mediated regulation of ATP biosynthesis as both a key player in the defense response against *E. faecalis* and possibly an exploit utilized by *S. enterica* during its pathogenesis. These findings indicate that NMUR-1 regulates energy homeostasis as part of defense against pathogen infection and provide valuable insights into the mechanism by which NMUR-1 regulates the specificity of innate immunity.

## RESULTS

### *S. enterica* and *E. faecalis* infections induce proteomic changes in *C. elegans*

To understand how *C. elegans* responds to *S. enterica* and *E. faecalis* infections at the protein level, we compared the proteomes of infected and uninfected WT N2 animals using a label-free quantitative proteomics approach. Five biological replicates of each experimental condition were analyzed with high resolution nano-HPLC tandem mass spectrometry. Quantifying the proteomes of WT animals exposed to *S. enterica* relative to control animals exposed to *E. coli* (worm food) yielded 2,959 proteins, of which 2,768 were identified with high confidence (1% false discovery rate, FDR). Among these proteins, 373 and 153 were significantly upregulated and downregulated at least 1.5-fold, respectively. Gene ontology (GO) analysis of the upregulated proteins revealed 64 significantly enriched biological processes, 26 of which are involved in the nematode’s energy production/electron transport chain (top 15 are shown in Fig. 1A). The most significantly upregulated group was “transmembrane transport” (q-value = 1.16E-19), with a notable subgroup being “ATP synthesis coupled proton transport” (q-value = 1.7E-8). These results indicate that infection with *S. enterica* caused a significant increase in transmembrane transport and in turn ATP synthesis in *C. elegans*. Indeed, *S. enterica* has been reported to cause an increase in reactive oxygen species (ROS) in the intestine of *C. elegans* as part of its pathogenesis (18), and a major source of ROS production in non-photosynthetic cells is as a by-product of ATP production in the mitochondria and as a product of oxidation of various biological molecules such as NADPH (19-21). Therefore, the increase in “ATP synthesis coupled proton transport” seen in our GO analysis is consistent with an increase in ATP production and induction of ROS production during *S. enterica* infection. We also performed GO analysis on the downregulated proteins and did not identify any significantly enriched processes.

**Figure 1.**
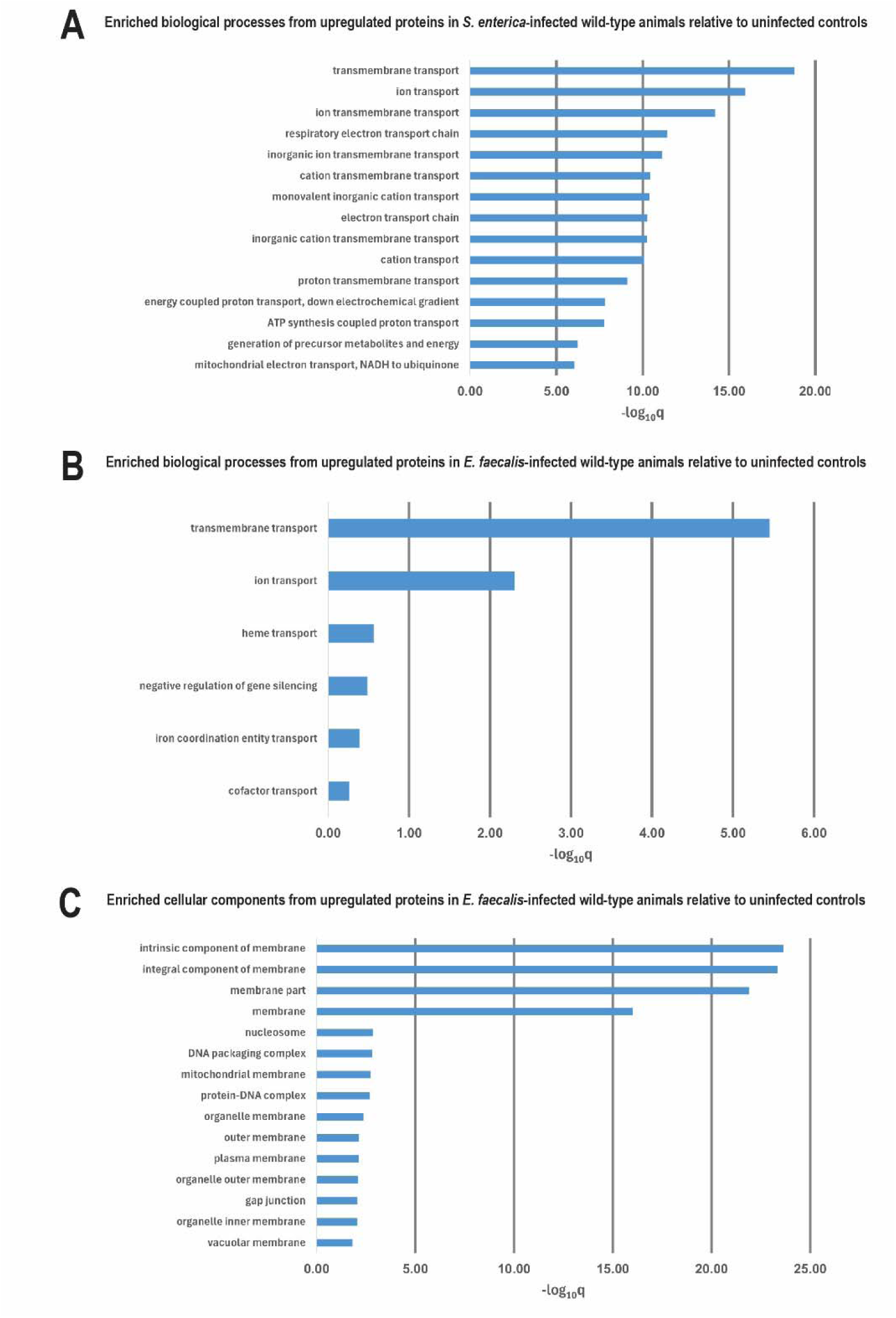
Infection changed the proteomes of WT animals. GO analyses were performed on upregulated proteins in WT animals infected with *S. enterica* **(A)** or *E. faecalis* **(B-C)** relative to uninfected control animals. The graphs show the top 15 most significantly enriched biological processes or cellular components in the infected samples. Bars represent the significance levels of enrichment expressed as -log_10_q values.

We next compared the proteomes of WT animals exposed to *E. faecalis* against control animals exposed to *E. coli* and identified 2,783 proteins with high confidence (1% FDR). Among these proteins, 240 and 120 were significantly upregulated and downregulated at least 1.5-fold, respectively. GO analysis of the upregulated proteins identified 6 significantly enriched biological processes (Fig. 1B). Similar to the *S. enterica*, *E. faecalis* infection increased the abundance of both ‘Transmembrane transport” (q-value = 3.54E-6) and “ion transport” (q-value = 4.93E-3) proteins. Further GO analysis of the proteins to identify their relevant cellular components revealed 24 enriched components, the two most significantly upregulated groups were “integral component of membrane” and “intrinsic components of membrane” (q-values = 4.65E-24 and 2.32E-24, respectively) (Fig. 1C). This increase in “integral” and “intrinsic” cell membrane components is consistent with a previous report of epithelial junction integrity being required for the defense against *E. faecalis* (22). GO analysis of the downregulated proteins during *E. faecalis* infection did not yield any significantly enriched biological processes. Overall, both *S. enterica* and *E. faecalis* infections triggered alternations in ATP biosynthesis and energy production in *C. elegans*, indicating important roles of energy homeostasis in the host response to infection. These results also demonstrate the effectiveness of our proteomic approach for detecting changes in protein expression during pathogen infection.

### NMUR-1 regulates the levels of ATP synthesis proteins

We previously showed that NMUR-1 suppresses worm survival against *S. enterica* but promotes worm survival against *E. faecalis* via transcriptional regulatory changes in the UPR and C-type lectin pathways, respectively (15). Here, we further investigated how the lack of NMUR-1 affected protein expression upon pathogen infection by comparing the proteomic changes of *nmur-1(ok1387)* mutant animals with those of WT animals when exposed to either *S. enterica* or *E. faecalis*.

Our proteomic data showed that during *S. enterica* infection, 74 proteins were upregulated, and 424 proteins were downregulated in *nmur-1(ok1387)* animals as compared to WT animals. GO enrichment analysis on the 74 upregulated proteins found a single enriched biological process, “pseudopodium” (q-value = 1.13E-3), the biological significance is unclear. A similar analysis of the 424 downregulated proteins detected 87 enriched biological processes, the most significant of which was “Transmembrane transport” (q-value = 3.46E-27) (Fig. 2A). This downregulation of transmembrane transport proteins is interesting, as it is juxtaposed to the proteome of WT animals on *S. enterica* where several of the same proteins were upregulated during infection. These results suggest that functional loss of NMUR-1 inhibits transmembrane transport and “ATP synthesis coupled proton transport”, which may contribute to the enhanced survival of *nmur-1(ok1387)* animals during *S. enterica* infection. As mentioned previously, ATP production is an important source of ROS (23-26). During *S. enterica* pathogenesis, the bacteria cause an increase in host ROS production, leading to cellular damage (18,27-29). Hindering ATP synthesis stifles ROS production, which, in turn, would slow *S. enterica* pathogenesis. This decrease in the abundance of ATP synthesis proteins and ROS in the *nmur-1(ok1387)* animals provides a possible mechanism responsible for their increased survival against *S. enterica*.

**Figure 2.**
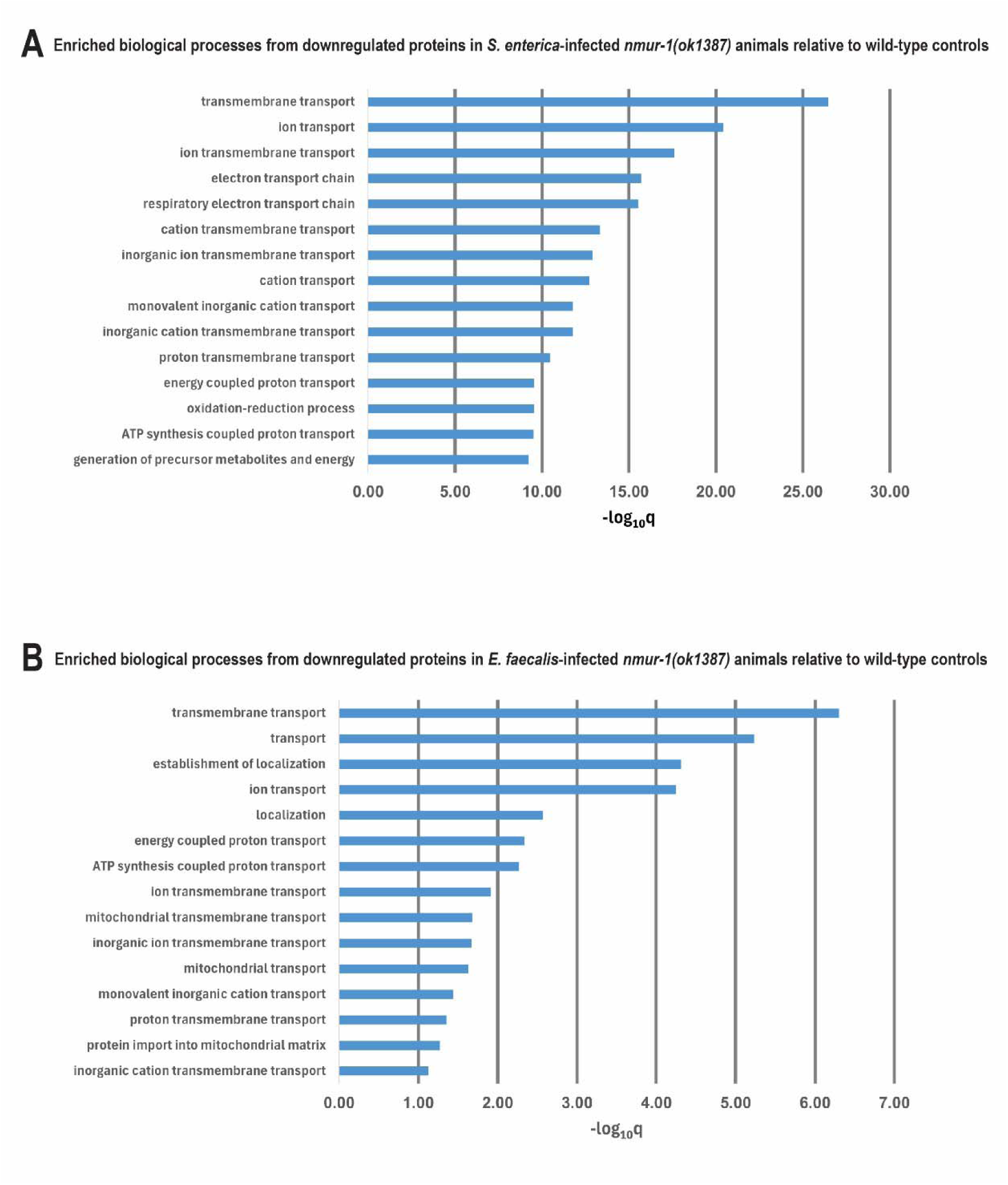
*nmur-1(ok1387)* mutant animals had a significant downregulation in ion transport and energy production proteins during infection. GO analyses were performed on downregulated proteins in *nmur-1(ok1387)* animals infected with *S. enterica* (A) or *E. faecalis* (B) relative to similarly infected WT control animals. The graphs show the top 15 most significantly enriched biological processes in mutant animals. Bars represent the significance levels of enrichment expressed as -log_10_q values.

We next compared the proteomes of *nmur-1(ok1387)* animals with those of WT animals during *E. faecalis* infection and found 85 and 221 proteins were upregulated and downregulated, respectively. While a GO enrichment analysis on the 85 upregulated proteins yielded no significantly enriched processes, a GO analysis on the 221 downregulated proteins identified 18 enriched biological processes. Again, the most significantly downregulated group was “Transmembrane transport” (q-value = 4.96E-7). Its subgroup “ATP synthesis coupled proton transport” was also found to be significantly downregulated (q-vaule = 5.38E-3) (Fig. 2B). How this downregulation of ATP synthesis-related proteins during *E. faecalis* infection affects survival of *nmur-1(ok1387)* animals is not yet clear. The reduced survival of *nmur-1(ok1387)* animals against *E. faecalis* is correlated to an increase in intestinal colonization by *E. faecalis* (15). To clear this intestinal colonization, *C. elegans* can excrete ROS from intestinal epithelium into the intestinal cavity. Previous reports indicate that the production of a specific ROS, H_2_O_2_, during *E. faecalis* infection is required for survival (30,31). During *E. faecalis* infection, *C. elegans* express BLI-3 that utilizes NADP(H) to generate H_2_O_2_ (30). However, generating NADP(H) is an ATP dependent process, and a reduction of available ATP would reduce the levels of NADPH and H_2_O_2_ (20,30,32-34). The lower abundance of ATP synthesis proteins and available ATP in *nmur-1(ok1387)* animals during *E. faecalis* infection could slow H_2_O_2_ production and lead to a reduced survival. Overall, we found that *nmur-1(ok1387)* mutant animals had a significantly lower expression of transmembrane transport and ATP synthesis proteins as compared to the WT control during both *S. enterica* and *E. faecalis* infection. A lack of ATP synthesis proteins may have contributed to the opposite survival phenotypes of *nmur-1(ok1387)* animals against these two pathogens through distinct mechanisms.

### ATP synthesis influences worm survival against both *S. enterica* and *E. faecalis* infections

To explore whether a lack of ATP synthesis proteins contributes to the survival phenotypes of *nmur-1(ok1387)* animals, we compared the proteins downregulated in both *S. enterica* and *E. faecalis* pathogenic conditions that fell within the same GO group “ATP synthesis coupled proton transport” to identify shared proteins. Seven proteins were identified as shared between the different conditions: ATP-3, -4, -5, ASB-1, -2, R53.4, and COX-5B. Five of the seven proteins were predicted subunits of the F_1_F_O_ ATP synthase complex: ATP-3, -4, -5, ASB-1, and -2. We narrowed our focus to ATP-3, -4, and -5 as these three proteins are subunits to the F_O_ peripheral stalk which acts as a stator for the F_1_ catalytic head and are homologs to human ATP5PD, ATP5PO, and ATP5PF, respectively (35-37).

To investigate whether disrupting the function of F_1_F_O_ ATP synthase has an influence over worm survival during *S. enterica* or *E. faecalis* infections, we treated the animals with *N,N’*-Dicyclohexylcarbodiimide (DCC) to chemically inhibit ATP synthesis. DCC is a classic inhibitor of ATP synthesis and functions by binding to the F_O_ subunit, inducing steric hindrance and preventing rotation of the F_O_ c-ring subunit required for ATP synthesis (38,39). WT animals exposed to DCC during *S. enterica* infection had a significant increase in survival as compared to the vehicle DMSO control (Fig. 3A). This is likely because the pathogenesis of *S. enterica* relies on an intrinsic induction of ROS in the intestinal epithelium of *C. elegans* (18), and a reduction in ATP synthesis leads to a reduction in ROS concentration, limiting the pathogenic tactic of *S. enterica.* However, the *nmur-1(ok1387)* mutants had no difference in survival against *S. enterica* between the DCC and vehicle control treatments (Fig. 3A), possibly due to the already downregulated key F_1_F_O_ ATP synthase subunits composing the peripheral stalk in the mutants (Fig. 2A). The peripheral stalk works in tandem with the membrane bound F_O_ c-ring to convert the flow of protons across the mitochondrial membrane into ATP (40,41). Without the peripheral stalk, ATP production by F_1_F_O_ ATP synthase is hindered and inhibition of the c-ring by DCC does not further inhibit production. Thus, the lack of key components forming the F_1_F_O_ ATP synthase peripheral stalk likely makes the *nmur-1(ok138)* mutant animals resistant to DCC treatment. Together, these results indicate that downregulation of F_1_F_O_ ATP synthase is responsible for the decrease in ATP production in *nmur-1(ok1387)* animals, which likely leads to an enhanced survival of *nmur-1(ok1387)* mutant animals against *S. enterica*.

**Figure 3.**
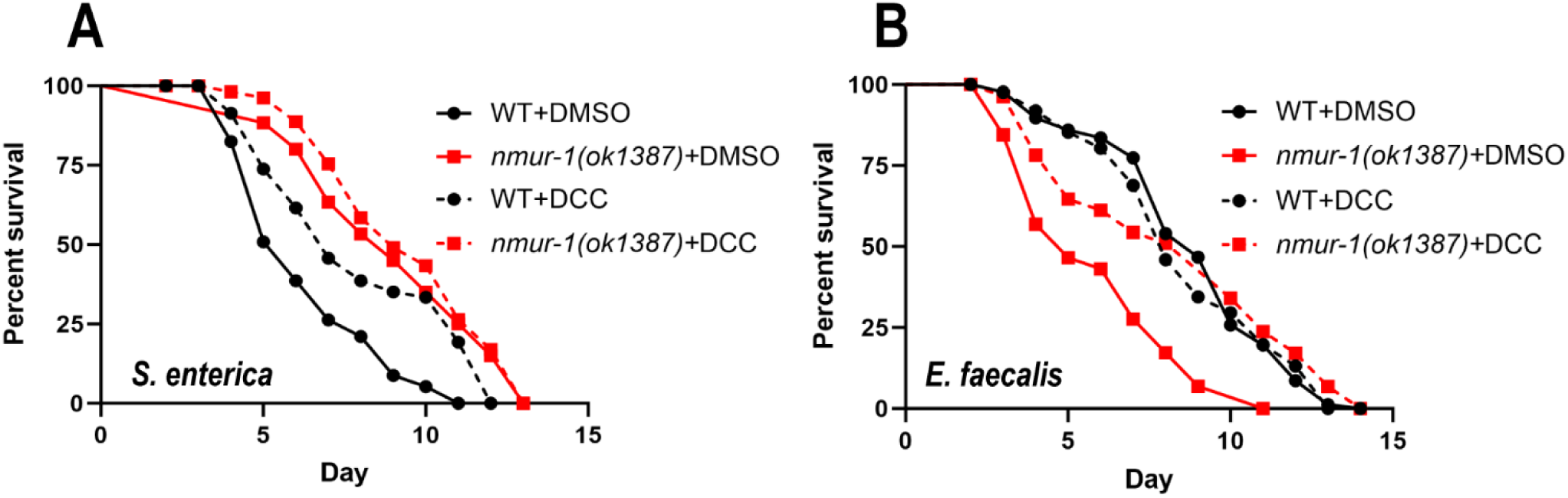
Chemical inhibition of F_1_F_O_ ATP synthase increased the survival of WT and *nmur-1(ok1387)* animals against *S. enterica* and *E. faecalis*, respectively. WT and *nmur-1(ok1387)* animals were exposed to *S. enterica* (A) or *E. faecalis* (B) on plates contain either DMSO or 20µM of DCC and scored for survival over time. Each graph is a representative of three independent experiments. Each experiment included *n* = 60 animals per strain. *p* values represent the significance levels of treatments relative to WT+DMSO: in **(A),** WT+DCC, *p* < 0.0001; *nmur-1(ok1387)+*DMSO, *p* < 0.0001; *nmur-1(ok1387)*+DCC, *p* < 0.0001; in **(B)**, WT+DCC, *p* = 0.3868; *nmur-1(ok1387)+*DMSO, *p* < 0.0001; *nmur-1(ok1387)*+DCC, *p* = 0.7616.

Exposure to DCC, however, significantly increased the survival of *nmur-1(ok1387)* mutants against *E. faecalis* relative to the vehicle control, but WT animals showed no significant difference between DCC treatment and the vehicle control (Fig. 3B). The increased survival in *nmur-1(ok1387)* animals with DCC treatment was unexpected and opposite to the above-described result with *S. enterica* infection. A possible explanation for this result is that DCC has antimicrobial properties as the chemical can inhibit both eukaryotic and prokaryotic F_1_F_O_ ATP synthase. When *E. faecalis* is exposed to DCC, the bacteria display a greater retention of the antibiotic gentamicin present in the media; this is because the exportation of gentamicin from cells requires ATP (42,43). Inhibition of ATP synthase by DCC increased the survival of *nmur-1(ok1387)* animals against *E. faecalis* because DCC compensates for the suppressed defense response in the mutant animals.

Next, we sought to determine if inactivation of individual ATP synthase subunits has any influence on worm survival against *S. enterica*. First, we crossed *atp-3(gk5653)* or *atp-5(gk5424)* knockout mutant animals with *nmur-1(ok1387)* animals to generate double mutants. We also silenced *atp-4* via RNA interference in both WT and *nmur-1(ok1387)* mutant animals (15). While inactivation of these genes displayed various survival phenotypes in WT animals (*atp-3(gk5653)* animals showed wild-type survival, and *atp-5(gk5424)* and atp-4 RNAi animals showed enhanced survival), *nmur-1(ok1387)* animals with such gene inactivation did not display any significant changes in survival time (Fig. 4, A and B), indicating that NMUR-1 and these ATP synthesis subunits might function in the same pathway to influence worm defense and survival against *S. enterica*.

**Figure 4.**
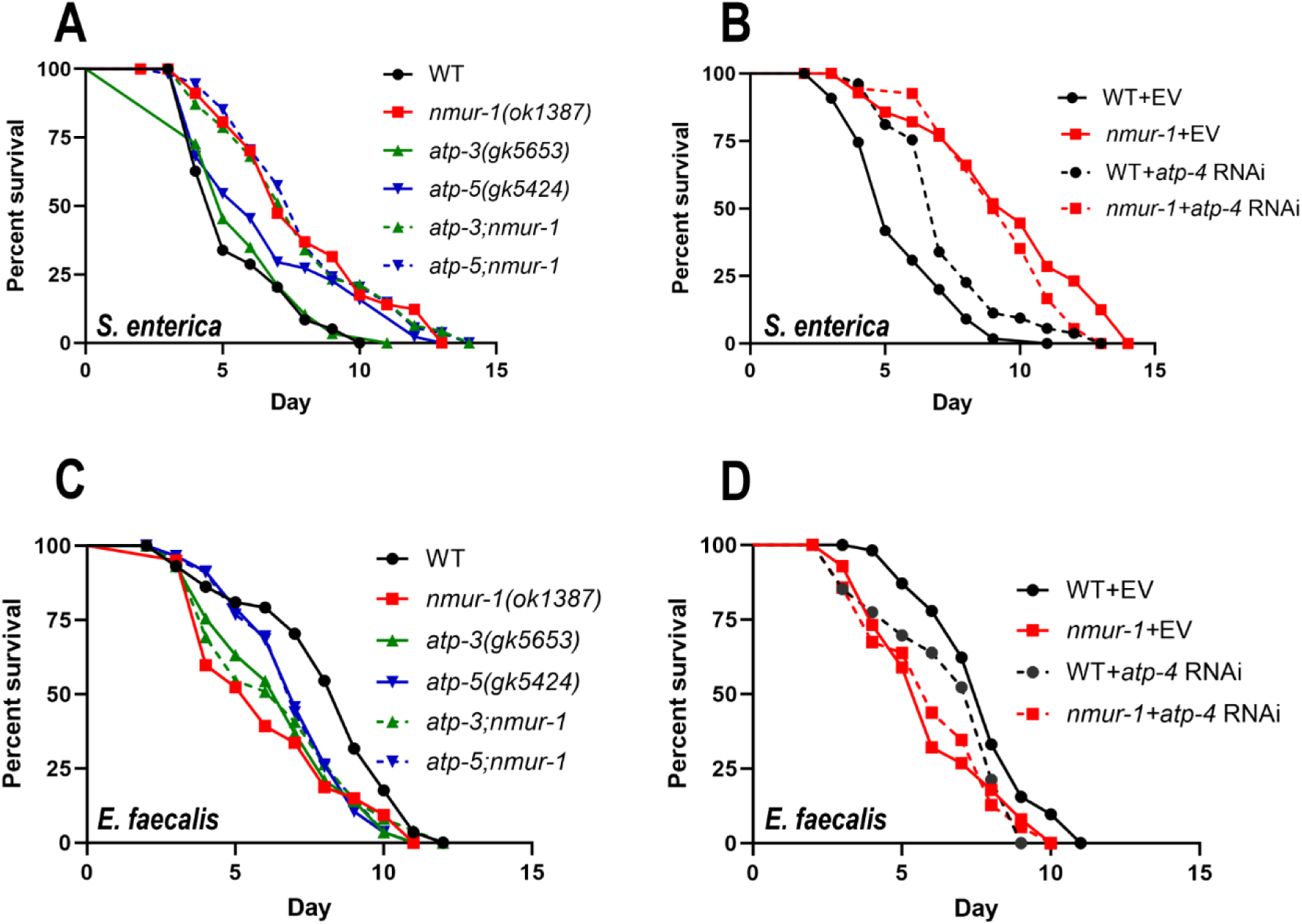
NMUR-1 regulated F_1_F_O_ ATP synthase subunits required for defense against S. enterica and E. faecalis. **(A)** WT, *nmur-1(ok1387), atp-3(gk5653), atp-5(gk5424), atp-3(gk5653);nmur-1(ok1387),* and *atp-5(gk5424);nmur-1(ok1387)* animals were exposed to *S. enterica* and scored for survival over time. The graphs are a representative of three independent replicates. Each experiment included *n* = 60 animals per strain. *p* values of mutants relative to WT: *nmur-1(ok1387), p* < 0.0001; *atp-3(gk5653)*, *p* = 0.8581; *atp-5(gk5424)*, *p* = 0.0053. *p* values of double mutants relative to *nmur-1(ok1387)*: *atp-3(gk5653)*;*nmur-1(ok1387)*, *p* = 0.8736; *atp-5(gk5424);nmur-1(ok1387)*, *p* = 0.9711. **(B)** WT and *nmur-1(ok1387)* animals grown on dsRNA for *atp-4* or the empty vector (EV) control were exposed to *S. enterica* and scored for survival over time. The graphs are a representative of three independent replicates. Each experiment included *n* = 60 animals per strain. *p* values of RNAi treatments relative to WT+EV: *nmur-1(ok1387)*+EV, *p* < 0.0001; WT+*atp-4*, *p* < 0.0001. *p* value of RNAi treatment relative to *nmur-1(ok1387)*+EV: *nmur-1(ok1387)*+*atp-4*, *p* = 0.0939**. (C)** WT, *nmur-1(ok1387), atp-3(gk5653), atp-5(gk5424), atp-3(gk5653);nmur-1(ok1387),* and *atp-5(gk5424);nmur-1(ok1387)* animals were exposed to *E. faecalis* and scored for survival over time. The graphs are a representative of three independent replicates. Each experiment included *n* = 60 animals per strain. *p* values of mutants relative to WT: *nmur-1(ok1387), p* < 0.0001; *atp-3(gk5653)*, *p* = 0.0002; *atp-5(gk5424)*, *p* = 0.0011. *p* values of double mutants relative to *nmur-1(ok1387)*: *atp-3(gk5653)*;*nmur-1(ok1387)*, *p* = 0.4323; *atp-5(gk5424);nmur-1(ok1387)*, *p* = 0.1924. **(D)** WT and *nmur-1(ok1387)* animals grown on dsRNA for *atp-4* or EV were exposed to *E. faecalis* and scored for survival over time. The graphs are a representative of three independent replicates. Each experiment included *n* = 60 animals per strain. *p* values of RNAi treatments relative to WT+EV: *nmur-1(ok1387)*+EV, *p* = 0.0001; WT+*atp-4*, *p* = 0.0113. *p* values of RNAi treatment relative to *nmur-1(ok1387)*+EV: *nmur-1(ok1387)*+*atp-4*, *p* = 0.8933.

We also examined the roles of individual ATP synthase subunits in worm survival against *E. faecalis*. Our results showed that *atp-3(gk5653)* and *atp-5(gk5424)* mutant animals displayed reduced survival against *E. faecalis* infection, however, *nmur-1(ok1387)* mutant animals carrying the same *atp-3* or *atp-5* mutation showed no significant difference in survival as compared to *nmur-1(ok1387*) single mutants (Fig. 4C). Silencing *atp-4* also significantly reduced the survival of WT animals against *E. faecalis* infection but did not significantly affect that of *nmur-1(ok1387)* animals (Fig. 4D). Taken together, while inactivation of *atp-3, atp-4*, or *atp-5* affected the survival of WT animals during *E. faecalis* infection, the survival of *nmur-1(ok1387)* mutant animals was unaffected by the loss of either of these genes during the same infection. These results suggest that the ATP synthesis pathway plays an important role in animal survival during both *S. enterica* and *E. faecalis* infection, and that *nmur-1(ok1387)* mutant animals suppress these proteins during infection.

### Functional loss of NMUR-1 reduces ATP concentrations during pathogen infection

Next, we examined the ATP concentrations in WT and *nmur-1(ok1387)* animals during *S. enterica* and *E. faecalis* infection using a luminescence method (44). Our results showed that during *S. enterica* infection, *nmur-1(ok1387)* mutant animals had significantly lower ATP concentrations compared to WT animals (Fig. 5A). Decreased ATP concentrations were also observed in *nmur-1(ok1387)* animals during *E. faecalis* infection (Fig. 5B). As a control, we quantified the ATP concentrations in WT animals exposed to either *S. enterica* or *E. faecalis* after being treated with DCC using the same method (DCC inhibits ATP synthesis and should reduce ATP concentrations). Comparing the ATP concentrations in animals exposed to either 20µM of DCC or the vehicle control, animals exposed to DCC had significantly reduced ATP concentrations after 24 hours (Fig 5). This validates our ATP quantification method and also confirms the efficiency of DCC inhibiting ATP production during a 24-hour pathogen exposure. Taken together, these results demonstrate that functional loss of NMUR-1 reduces ATP concentrations during pathogen infection, which may contribute to the survival phenotypes of *nmur-1(ok1387)* animals against *S. enterica* and *E. faecalis*.

**Figure 5.**
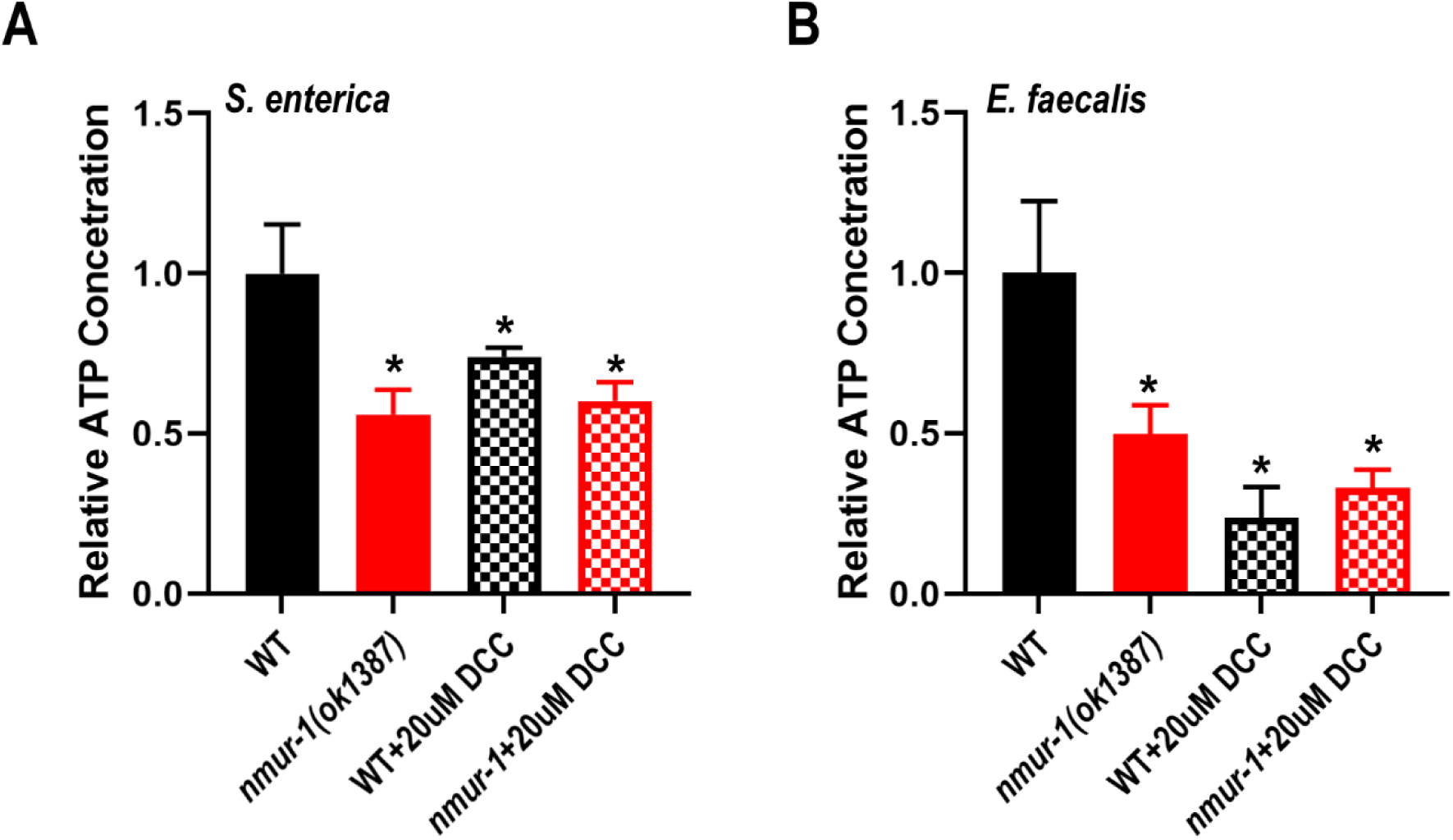
NMUR-1 regulated ATP levels during pathogen infection. WT and *nmur-1(ok1387)* animals were exposed to *S. enterica* **(A)** or *E. faecalis* **(B)** for 24 hours with or without 20µM DCC followed by ATP concentration measurements. The graphs show combined results of three independent experiments. In each experiment, 30 animals of each strain under either condition were used. Error bars represent standard deviation. Asterisk (*) denotes a significant difference (*p* < 0.05) between treated animals and untreated WT controls, as analyzed using Dunnett’s multiple comparison test.

### Functional loss of NMUR-1 reduces ROS concentrations during pathogen infection

Next, we sought to determine if ROS concentrations were also affected by the suppression of ATP biosynthesis in *nmur-1(ok1387)* mutant animals. To determine the ROS concentrations of animals infected with either *S. enterica* or *E. faecalis*, we followed a previously established protocol utilizing Dichlorodihydrofluorescein Diacetate (45). WT and *nmur-1(ok1387)* mutant animals were exposed to *S. enterica* or *E. faecalis* for 24 hours prior to ROS quantification. Compared to WT animals during either *S. enterica* or *E. faecalis* infection, *nmur-1(ok1387)* mutant animals had a significantly lower fluorescent intensity correlating to a lower ROS concentration (Fig. 6). These findings support our previous notion that lower ATP biosynthesis may lead to lower ROS concentrations in *nmur-1(ok1387)* mutant animals. Overall, these results are consistent with previous studies highlighting the importance of ROSs during *S. enterica* and *E. faecalis* infection, and *nmur-1* plays a significant role in regulating its production.

**Figure 6.**
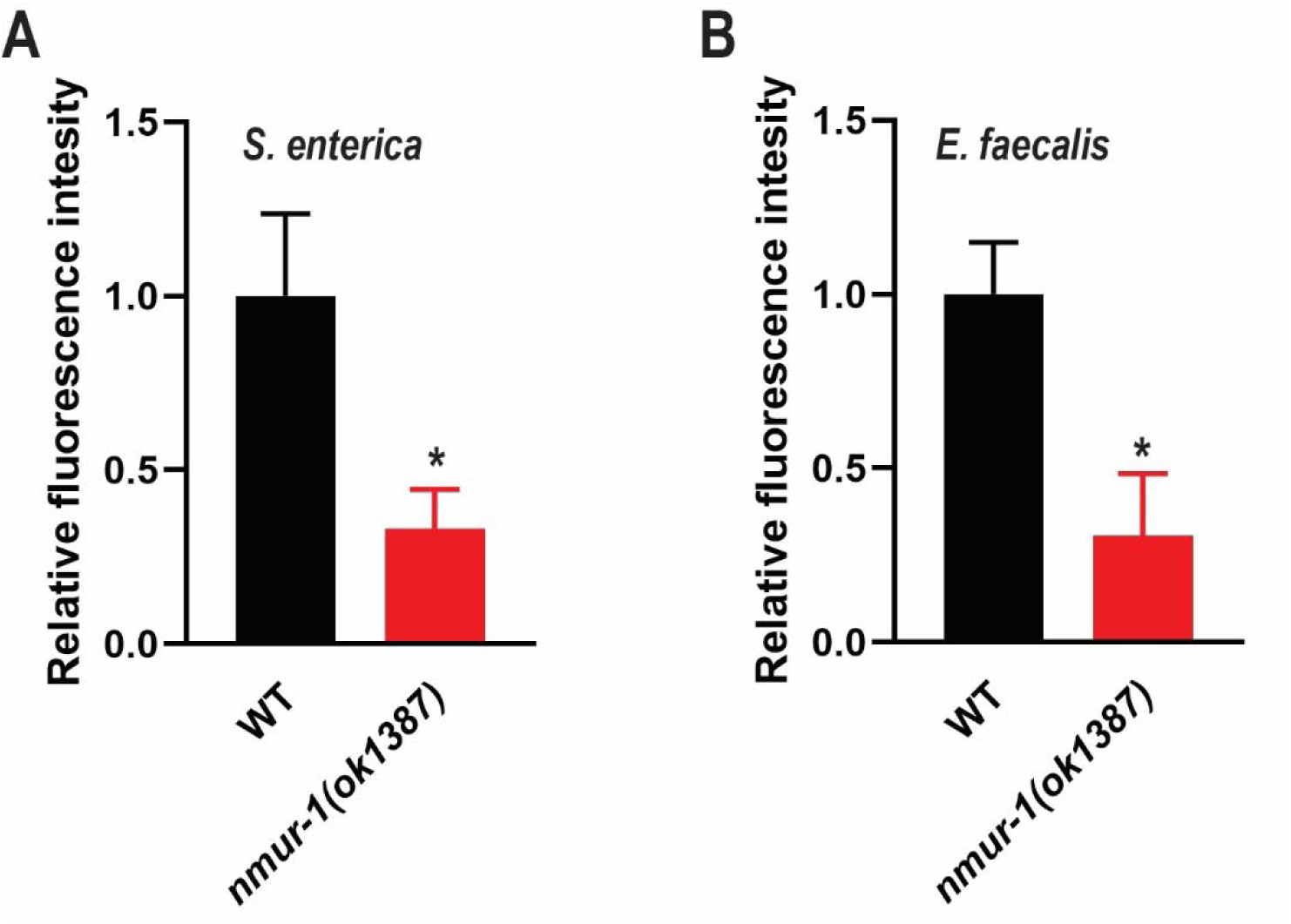
ROS concentrations were lower in *nmur-1(ok1387)* animals than in WT animals exposed to pathogens. WT and *nmur-1(ok1387)* animals were exposed to *S. enterica* **(A)** or *E. faecalis* **(B)** for 24 hours. The animals were stained for ROS production using H_2_DCFDA and the resulting GFP intensity was measured using a Filtermax 5 plate reader. The graphs show combined results of three independent replicate experiments which measured the ROS production of 30 animals per replicate. The results are relative to the normalized fluorescence in untreated WT animals, and asterisk (*) denotes a significant difference (*p* < 0.05) as determined by Student’s t-test.

### The NMUR-1 endogenous ligand CAPA-1 and its expressing neurons ASG regulate ATP biosynthesis during *E. faecalis* infection

Our previous research revealed that the NMUR-1 endogenous ligand CAPA-1 expressed by the ASG neurons was required for the survival of *C. elegans* against *E. faecalis* but dispensable in defense against *S. enterica* (15). Here, we focused on investigating whether ATP biosynthesis is regulated by the NMUR-1/CAPA-1/ASG signaling in the context of *E. faecalis* infection. The knockout mutant *capa-1(ok3065)* displays a reduced survival phenotype against *E. faecalis* similar to *nmur-1(ok1387)* (15) (Fig. S1). Since CAPA-1 is only expressed in a pair of ASG amphid neurons (46), we questioned if removal of these neurons would also have a similar effect on survival against *E. faecalis*. Measuring the survival of an ASG-ablated strain against *E. faecalis,* we found the lack of ASG neurons indeed led to reduced survival (Fig. 7A).

**Figure 7.**
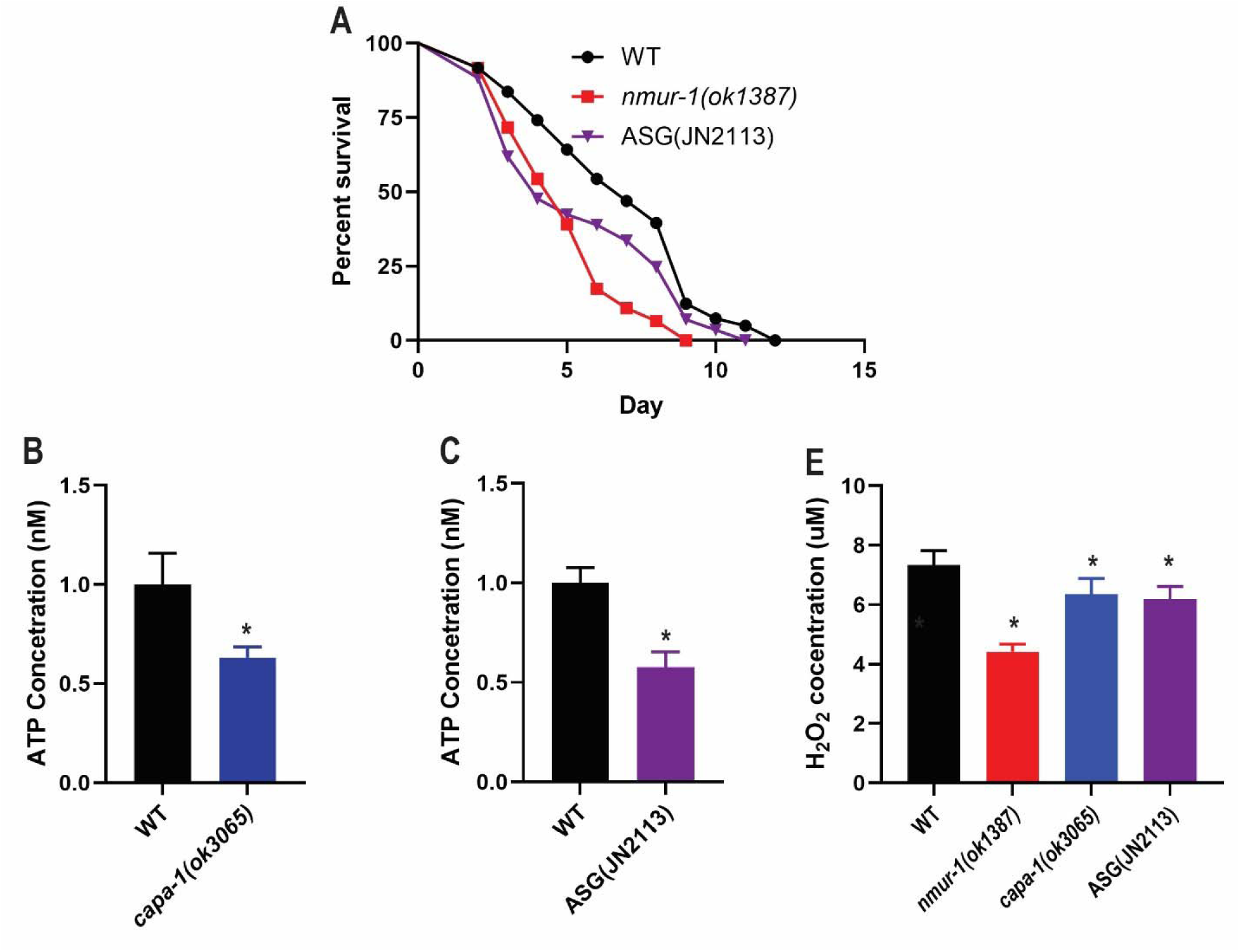
Loss of CAPA-1 or ASG neurons partially mimicked the phenotypes of *nmur-1(ok1387)* animals against *E. faecalis*. **(A)** WT, *nmur-1(ok1387)*, and ASG ablated JN2113 animals were exposed to *E. faecalis* and scored for survival over time. The graphs are a representative of three independent replicates. Each experiment included *n* = 60 animals per strain. *p* values represent the significance levels of mutants relative to WT: *nmur-1(ok1387), p* < 0.0001; JN2113, *p* = 0.0292. **(B-C)** WT, *capa-1(ok3065*, and JN2113 animals were exposed to *E. faecalis* for 24 hours followed by ATP concentration measurements. The graphs show combined results of three independent experiments. In each experiment, 30 animals of each strain were used. Error bars represent standard deviation. Asterisk (*) denotes a significant difference (*p* < 0.05) between mutants and WT animals. **(D)** Animals were exposed to *E. faecalis* for 24 hours. H_2_O_2_ concentrations were quantified using Amplex Red fluorescent dye after a 30-minute incubation. The graphs show combined results of three independent replicates which measured the respiration of 30 animals per replicate. The results were normalized to the number of animals collected per experiment. Error bars represent standard deviation. Asterisk (*) denotes a significant difference (*p* < 0.05) between treatment and the untreated WT control, as determined by Dunnett’s multiple comparison test.

Quantifying ATP concentrations in both *capa-1(ok3065)* knockout animals and ASG-ablated animals during *E. faecalis* infection found both strains also had significantly reduced ATP levels as compared to WT animals (Fig. 7C and 7D). These results indicate that the CAPA-1/NMUR-1/ASG signaling regulates ATP biosynthesis in the context of *E. faecalis* infection, which likely contributes to the overall defense against this pathogen.

## DISCUSSION

In our current study, we have shown that *C. elegans* lacking the neuronal GPCR NMUR-1 have an altered proteome when challenged with the pathogen *S. enterica* or *E. faecalis*. We focused our attention on these two pathogens as *nmur-1* mutant animals display opposing survival phenotypes when infected with these pathogens, which can serve as a useful model for understanding the roles of NMUR-1 in mediating the specificity of innate immune responses (15). Utilizing quantitative mass spectrometry, we found that animals with a *nmur-1* knockout have a significant decrease in expression of proteins responsible for ATP biosynthesis during infection. This is interesting as studies have already pointed to the role of ATP biosynthesis and the mitochondria in regulating the innate and adaptive immune responses (47-49). We hypothesized that this decrease in ATP biosynthesis plays a role in the survival phenotypes of *nmur-1(ok1387)* mutant animals challenged with *S. enterica* and *E. faecalis*. Indeed, inhibition of the ATP synthesis caused WT animals to partially mimic the survival of *nmur-1(ok1387)* mutant animals during *S. enterica* or *E. faecalis* infection. We also questioned if specific neurons played any role in regulating ATP production through an NMUR-1-dependent pathway. Our previous studies have shown that the endogenous ligand CAPA-1 was required for survival against *E. faecalis* and other labs have shown that CAPA-1 is expressed in the ASG neurons (15,46). Indeed, we found that CAPA-1 and the ASG neurons are required for *C. elegans* survival against *E. faecalis* and a lack of either causes a decrease in ATP production (Fig. 7).

How is NMUR-1, a single neuronal GPCR, able to regulate ATP biosynthesis in response to various pathogens but mediate differing survival phenotypes? During *E. faecalis* infection, ASG neurons secrete CAPA-1 that binds to NMUR-1 and promotes ATP and ROS production. This signaling cascade promotes the survival of *C. elegans* against *E. faecalis*. In the case of *S. enterica* infection, however, animals lacking CAPA-1 do not share an enhanced resistant phenotype like NMUR-1 knockout animals (15). This points to the possibility that upon *S. enterica* infection, *C. elegans* is not activating NMUR-1 via CAPA-1 and triggering the downstream increase in ATP and ROS synthesis. Instead, *S. enterica* may produce an exogenous ligand that binds to NMUR-1. This is supported by another study investigating how *S. enterica* is able to colonize the intestine of *C. elegans* (18). The authors found that animals infected with *S. enterica* had an increased concentration of ROS during infection, and that this increase in ROS was necessary for killing by *S. enterica* (18). Without NMUR-1, *S. enterica* is unable to trick *C. elegans* into overproducing ROSs, causing the mutant animals to have an enhanced resistance to the pathogen. This dual role as both a receptor for mounting a proper immune response against *E. faecalis* and a vulnerability against *S. enterica* places NMUR-1 in a pivotable space in host-pathogen interactions. Due to its roles in host-pathogen interactions and the highly conserved nature of NMUR-1 across multiple species, future work studying the mechanisms and signaling pathways of NMUR-1 during pathogen infection would provide valuable insights to the communication between the nervous system and other nonneuronal tissues under pathogenic and non-pathogenic conditions.

## MATERIALS AND METHODS

### C. elegans strains

The following *C. elegans* strains were maintained as hermaphrodites at 20°C, grown on modified Nematode Growth Media (NGM) (0.35% instead of 0.25% peptone) with 15µg/mL tetracycline, and fed *E. coli* HT115 (50). The wild-type animal strain was *C. elegans* Bristol *N2*. The *atp-3(gk5653)* and *atp-5(gk5424)* were obtained from the *Caenorhabditis* Genetics Center (University of Minnesota, Minneapolis, MN). The *nmur-1(ok1387)* animals were from a previous study performed by our lab. All mutant animals were backcrossed with wild-type Bristol *N2* animals at least three times before experimental use. All genotypes were confirmed using PCR or visual inspection of GFP expression in the pharyngeal bulb for *atp-3(gk5653)* and *atp-5(gk5424)*. All animals used for this study were synchronized 65-hour old young adults unless otherwise noted in the specific methodology.

### Bacterial strains

The following bacteria strains were grown using standard conditions (51). *Escherichia coli* strain HT115, *Salmonella enterica* strain SL1344. *Enterococcus faecalis* strain OG1RF. E. coil HT115 and S. enterica SL1344 were cultured in Luria-Bertani (LB) broth and agar which was prepared following the manufacturer’s recommendations. All assays for *E. coil* HT115 and *S. enterica* were performed on NGM. *E. faecalis* OG1RF was cultured in brain heart infusion (BHI) broth and agar which was prepared following the manufacturer’s recommendations. All assay using *E. faecalis* OG1RF were performed on BHI media containing 10µg/mL of gentamycin.

### *C. elegans* pathogen exposure and protein collection

Gravid adult wild-type and *nmur-1(ok1387)* animals were lysed using a solution of sodium hydroxide and bleach (volume ratio 5:2), washed using M9 buffer, and eggs were synchronized for 22 hours in S-basal liquid medium at room temperature. Synchronized L1 larval animals were transferred onto modified NGM plates seeded with *E. coli* HT115 and grown at 20°C for 48 hours until the animals had reached the L4 larval stage. The synchronized wild-type and *nmur-1(ok1387)* L4 larval stage animals were collected and transferred to plates seeded with either *S. enterica* SL1344, *E. faecalis* OG1RF, or *E. coli* HT115 for 24 hours at 25°C. Infected and uninfected controls (exposed to *E. coli* HT115) were collected and washed 5 times in M9 buffer containing a 1x concentration of protease inhibitors (Halt-protease inhibitor cocktail, ThermoFisher Sci.) to remove any bound bacteria. After the last wash and removal of the supernatant, each sample was snap frozen in a bath of ethanol and dry ice. Five biological replicates of *S. enterica*, *E. faecalis*, and *E. coli* treated animals were collected, respectively. The frozen samples were submitted to the Tissue Imaging and Proteomics Laboratory at Washington State University (Pullman, WA) for mass spectrometry-based quantitative proteomics analyses.

### Protein sample preparation

The worm pellet was freeze dried and ground into fine powder using a single 2.8 mm i.d. steel ball with a TissueLyser II (Qiagen, Valencia, CA) at a frequency of 30 Hz for 30 sec, followed by the addition of 100 µL extraction solvent of PBS buffer, pH 7.5, with protease inhibitor and vortexing. Supernatants were collected after centrifugation at 16,000 × g (10 min, 4 °C). Protein was then quantified with a Qubit Protein Assay Kit (Life Technologies, Carlsbad, CA) in compliance with the manufacturer’s protocol. Disulfide bonds were reduced using 100 mM dithiothreitol (DTT) at a ratio of 1:10 DTT/sample volume and incubated at 50 °C for 45 min. Cysteine bonds were then alkylated with 200 mM iodoacetamide at the same volume ratio for 20 min at room temperature. Finally, protein was digested with trypsin (G-Biosciences, St. Louis, MO) at a 1:50 ratio of trypsin/protein, and incubated at 37 °C for 12 hours.

### High resolution nano-HPLC tandem mass spectrometry analysis

The peptide samples were subjected to Thermo Scientific Orbitrap Fusion Tribrid with an Easy-nLC 1000 ultra-high pressure LC on a Thermo Scientific PepMap 100 C18 column (2□μm, 50□μm□×□15□cm). The peptides were separated over 115□min gradient eluted at 400□nL/min with 0.1% formic acid (FA) in water (solvent A) and 0.1% FA in acetonitrile (solvent B) (5–30% B in 85□min, followed by 30–50% B over 10□min and 50–97% B over 10□min). The run was completed by holding a 97% B for 10□min. MS1 data was acquired on an Orbitrap Fusion mass spectrometry using a full scan method according to the following parameters: scan range 400–1500□m/z, Orbitrap resolution 120,000; AGC target 400,000; and maximum injection time of 50□ms. MS2 data were collected using the following parameters: rapid scan rate, HCD collision energy 35%, 1.6□m/z isolation window, AGC 2,000 and maximum injection time of 50□ms. MS2 precursors were selected for a 3□s cycle. The precursors with an assigned monoisotopic m/z and a charge state of 2–7 was interrogated. The precursors were filtered using a 60□s dynamic exclusion window. MS/MS spectra were searched using Thermo Scientific Proteome Discoverer software version 2.0 with SEQUEST® against uniprot *Caenorhabditis elegans* database (TaxID□=□6239). Precursor and fragment mass tolerances were set to 10□ppm and 0.8□Da respectively and allowing up to two missed cleavages. The static modification used was carbamidomethylation (C). The high confidence level filter with false discovery rate (FDR) of 1% was applied to the peptides. Protein relative quantitation was achieved by extracting peptide areas with the Proteome Discoverer 2.0 (Thermo Scientific, San Jose, CA) and 3 unique peptides per protein were used for the protein quantitation analysis. The mass spectrometry proteomics data have been deposited to the ProteomeXchange Consortium via the PRIDE65 partner repository with the dataset identifier PXD045826.

### Survival assay

Wild-type and mutant animals were synchronized by egg-laying. Briefly, well-fed gravid adult animals were transferred to fresh *E. coli* HT115 seeded NGM plates and incubated for 45 minutes at 25°C. Adult animals were removed after 45 minutes, and the synchronized offspring were grown at 20°C for 65 hours to reach young adult stage. Bacterial lawns for the survival assays were prepared by culturing pathogenic bacteria for 15∼16 hours in either LB broth for *S. enterica* or BHI broth for *E. faecalis* OG1RF at 37°C in a shaking incubator. A 30µL drop of the fresh bacterial culture was placed on 3.5cm plates of either modified NGM or BHI media with 10 µg/mL of gentamicin. Plates were incubated at 37°C for 15∼16 hours, cooled to room temperature, and then seeded with synchronized 65-hour-old young adult animals. The survival assays were performed at 25°C, and live animals were transferred daily to fresh plates until egg laying ceased. Animals were scored at the times indicated and were considered dead when they failed to respond to touch.

### RNA interference

RNAi was conducted using the Ahringer group library and feeding synchronized L4 larval *C. elegans E. coli* strain HT115(DE3) expressing double-stranded RNA (dsRNA) that was homologous to a target gene. (52,53). Before exposure, all RNAi clone plasmids were isolated, digested with KpnI, and Sanger sequenced using a T7 promoter primer to check for gene specificity. *E. coli* with the appropriate dsRNA vector were grown in LB broth containing ampicillin (100µg/mL) at 37°C for 15∼16 hours and 120µL was plated on modified NGM plates containing 100µg/mL ampicillin and 3mM isopropyl β-D-thiogalactoside (IPTG). The bacteria were allowed to grow for 15∼16 hours at 37°C. The plates were cooled away from direct light before the synchronized L3 larval animals were placed on the bacteria. Synchronized L4 larvae worms were placed on RNAi or vector control plates for 24 hours at 20 °C. Adults were then transferred to survival assay plates. Clone identity was confirmed by sequencing at Eton Bioscience Inc. *unc-22* RNAi was included as a positive control in all experiments to account for RNAi efficiency.

### ATP Quantification

Synchronized wild-type and *nmur-1(ok1387)* animals and bacteria plates were prepared as described in survival assay. 65-hour-old animals were placed on either *S. enterica* SL1344 or *E. faecalis* OG1RF plates. The animals were incubated at 25°C for 24 hours. Following incubation, animals were randomly picked from the bacteria plates, placed into 50µL of M9 buffer and washed three times then snap frozen in a bath of dry ice and ethanol. Suspended animals were boiled for 15 mins at 95°C then placed on ice for 5 mins. Supernatant was collected and transferred to a new 1.5mL tube after centrifugation at 14,800 x *g* for 10 mins at 4°C. ATP quantification was performed using the ATP Determination kit by ThermoFisher (A22066) following the manufacturer’s protocol using 10µL of 10-fold diluted supernatant in a 100µL reaction. Luminescence was recorded using a Molecular Devices FilterMax F5.

### *N,N’*-Dicyclohexylcarbodiimide treatment

NGM and BHI media were prepared following the method described above with the addition of either DMSO or 20µM of *N,N’*-Dicyclohexylcarbodiimide (DCC). A 20mM solution of DCC was prepared by dissolving 41.266mg of DCC into 10mL of DMSO. The solution was prepared fresh and not allowed to be stored for more than one day as the combination of DCC and DMSO causes the DCC to undergo Pfitzer-Moffatt oxidation and precipitate out of solution. 1mL of DMSO or 1mL 20mM of DCC in DMSO was added to 1L of media before it solidified. Animals were synchronized as described above in the absence of DCC. Synchronized animals were moved to DCC containing NGM media plates 24 hours before pathogen exposure. The survival assays were carried out in the same manner as described above.

### H_2_O_2_ respiration quantification

H_2_O_2_ respiration was quantified following a previous describe protocol (54). In short, synchronized wild-type and *nmur-1(ok1387)* animals and bacteria plates were prepared using the bleach syncing method as described above. 65-hour-old animals were collected in M9 buffer and placed on *E. faecalis* OG1RF plates then placed in a 25°C incubator for 24 hours. After 24 hours, the animals were washed off the bacteria plates using M9 buffer, the adult animals were allowed to settle to the bottom of a 15mL conical tube before the supernatant was removed. The animals were washed twice to remove excess bacteria and L1 larval animals. The animal pellet was then washed twice with 5mL of Amplex assay reaction buffer to further remove excess bacteria before being transferred to an empty BHI plate. 30 washed adult animals were transferred from the BHI plate without the use of bacteria as an adhesive to a single well of a black clear bottom 96-well plate filled with 50µL of 1x Amplex assay reaction buffer. 50µL of Amplex working solution was then added to each well, the final concentration of reagents were 50µM of Amplex Red and 0.1U/mL of horseradish peroxidase. Animals were placed on the opposite end of the assay plate away for the standards to avoid cross contamination. The assay was measured over the course of 2 hours using a Molecular Devices FilterMax F5, measurements were taken every 15 minutes.

### ROS quantification using 2’,7’-Dichlorodihydrofluorescein Diacetate

ROS quantification using 2’,7’-Dichlorodihydrofluorescein Diacetate (H_2_DCFDA) was performed following a previously described protocol (45). In short, synchronized wild-type and *nmur-1(ok1387)* animals and bacteria plates were prepared using the egg-laying method as described above. 65-hour-old animals were collected in M9 buffer and were placed on either *S. enterica* SL1344 or *E. faecalis* OG1RF plates, the plates were then placed in a 25°C incubator for 24 hours. After 24 hours, the animals were transferred to an unseeded NGM plate and allowed to crawl to separate the adults from eggs and L1 larval animals. 30 adult animals were transferred to a single well of a black clear bottom 96-well plate filled with 50µL of M9 buffer. 50µL of 50µM H_2_DCFDA was pipetted into each well for a final concentration of 25µM. The assay was measured over the course of 6 hours using a Molecular Devices FilterMax F5, measurements were taken every 30 minutes.

### Quantification and statistical analysis

Survival curves were plotted using GraphPad PRISM (version 10) computer software. Survival was considered different from the appropriate control indicated in the main text when P < 0.05. PRISM uses the product limit or Kaplan-Meier method to calculate survival fractions and the log-rank test, which is equivalent to the Mantel-Haenszel test, to compare survival curves, and qRT-PCR results were analyzed using two-sample t-tests for independent samples; P-values < 0.05 are considered significant. All experiments were repeated at least three times, unless otherwise indicated. Statistical details for each figure are listed in its corresponding figure legend.

## ACKNOWLEDGEMENTS

We thank Drs. Jennifer Watts, Kathryn Meier, and Mike Gibson at Washington State University for constructive discussions. We thank Dr. Anna Berim at the Tissue Imaging, Metabolomics and Proteomics Laboratory (TIMPL), WSU for depositing the mass spectrometry proteomics data to the ProteomeXchange Consortium. Some worm strains in this study were provided by the Caenorhabditis Genetics Center, which is funded by the NIH Office of Research Infrastructure Programs (P40 OD010440). This work was supported by NIH (R35GM124678 to J.S.) and the Department of Translational Medicine and Physiology, Elson S. Floyd College of Medicine, WSU-Spokane (startup to Y.L.). The funders had no role in study design, data collection and interpretation, or the decision to submit the work for publication.

## AUTHOR CONTRIBUTIONS

P.W. designed and performed experiments and analyzed data. D.B. performed mass spectrometry proteomics experiments and analyzed data. Y.L., K.R., and J.S. designed experiments and analyzed data. D.G. designed experiments. P.W., Y.L., and J.S. wrote the paper.

## MATERIAL AVAILABILITY

The *C. elegans* strains and recombinant DNA generated in this study will be shared upon request, but we may require payment to cover shipment and completion of a Material Transfer Agreement for possible commercial applications.

## DATA AND CODE AVAILABILITY

- The raw proteomic data was deposited to the Proteome exchange via Proteomics IDEntifications database (PRIDE). All of this data can be accessed through PRIDE with the accession number PXD045826.
- No original code was created during the course of this study.
- Any additional information required to reanalyze the data reported in this paper is available from the lead contact upon request.

---

**Figure S1.**
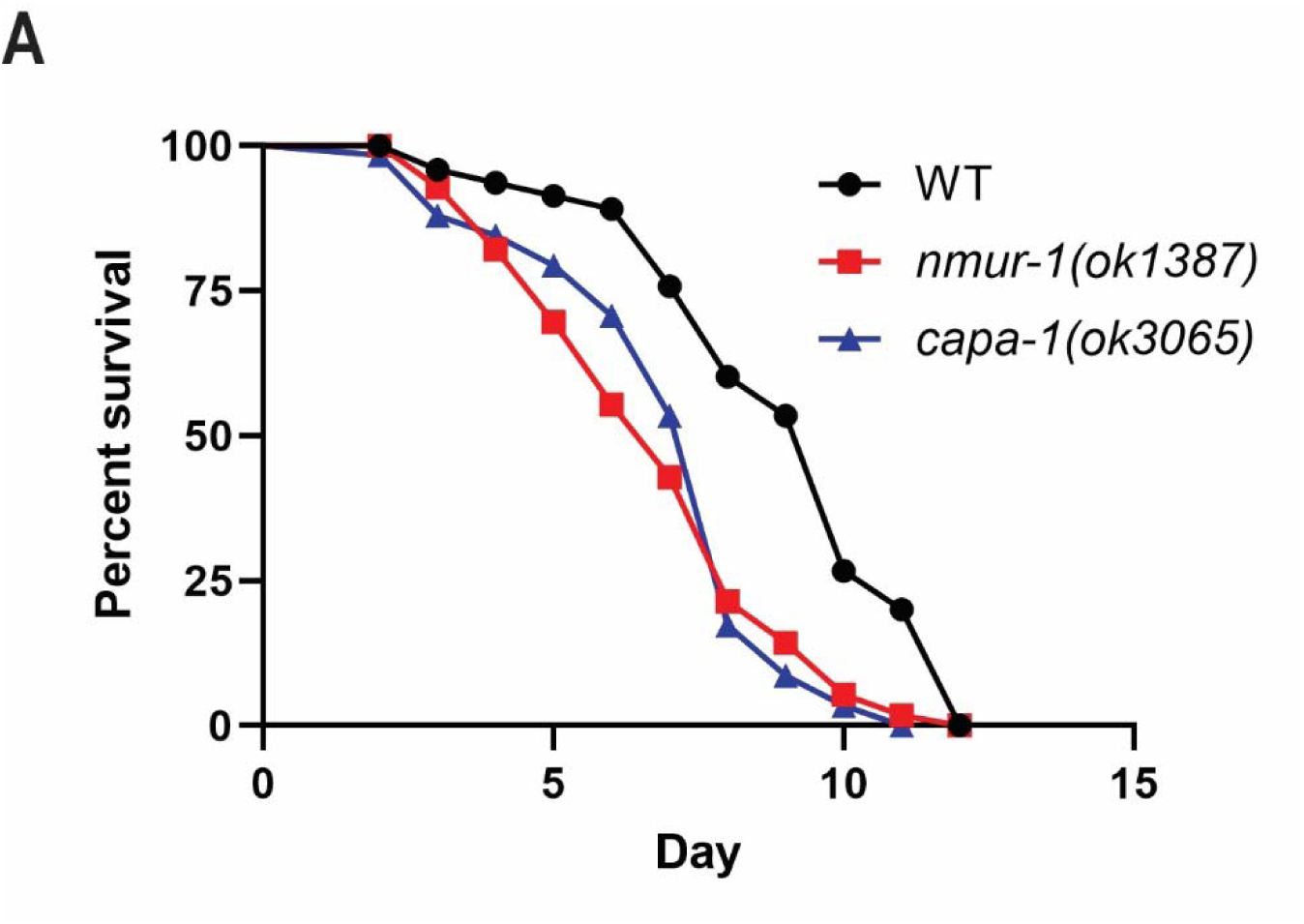
Loss of CAPA-1 reduced *C. elegans* survival against *E. faecalis*, mimicking the survival phenotype of *nmur-1(ok1378)* animals. WT, *nmur-1(ok1387)*, and *capa-1(ok3065)* animals were exposed to *E. faecalis* and scored for survival over time. The graphs show a representative of three independent replicates. Each experiment included *n* = 60 animals per strain. *p* values represent the significance levels of mutants relative to WT: *nmur-1(ok1387)*, *p* < 0.0001; *capa-1(ok3065)*, *p* < 0.0001.

